# Single-Cell Transcriptomic Analysis of Kaposi Sarcoma

**DOI:** 10.1101/2024.05.01.592010

**Authors:** D. A. Rauch, P. Valiño Ramos, M. Khanfar, J. Harding, A. Joseph, O Griffith, M Griffith, L. Ratner

## Abstract

Kaposi Sarcoma (KS) is a complex tumor caused by KS-associated herpesvirus 8 (KSHV). Histological analysis reveals a mixture of “spindle cells”, vascular-like spaces, extravasated erythrocytes, and immune cells. In order to elucidate the infected and uninfected cell types in KS tumors, we examined skin and blood samples from twelve subjects by single cell RNA sequence analyses. Two populations of KSHV-infected cells were identified, one of which represented a proliferative fraction of lymphatic endothelial cells, and the second represented an angiogenic population of vascular endothelial tip cells. Both infected clusters contained cells expressing lytic and latent KSHV genes. Novel cellular biomarkers were identified in the KSHV infected cells, including the sodium channel SCN9A. The number of KSHV positive tumor cells was found to be in the 6% range in HIV-associated KS, correlated inversely with tumor-infiltrating immune cells, and was reduced in biopsies from HIV-negative individuals. T-cell receptor clones were expanded in KS tumors and blood, although in differing magnitudes. Changes in cellular composition in KS tumors were identified in subjects treated with antiretroviral therapy alone, or immunotherapy. These studies demonstrate the feasibility of single cell analyses to identify prognostic and predictive biomarkers.

**Author Summary:** Kaposi sarcoma (KS) is a malignancy caused by the KS-associated herpesvirus (KSHV) that causes skin lesions, and may also be found in lymph nodes, lungs, gastrointestinal tract, and other organs in immunosuppressed individuals more commonly than immunocompetent subjects. The current study examined gene expression in single cells from the tumor and blood of these subjects, and identified the characteristics of the complex mixtures of cells in the tumor. This method also identified differences in KSHV gene expression in different cell types and associated cellular genes expressed in KSHV infected cells. In addition, changes in the cellular composition could be elucidated with therapeutic interventions.

## Introduction

Kaposi sarcoma-associated herpesvirus (KSHV), which is also known as human gammaherpesvirus 8 (HHV-8), is a member of the rhadinovirus genus, and was first identified in 1994 (1). The highly-conserved, circular, 165-kb double stranded DNA genome of KSHV has a 140-kb unique region encoding ∼90 genes flanked by 20-30 kb of terminal repeat sequences (2). The viral genome is maintained as an episome in infected cells and persists in a latent state during which it expresses a latency-associated nuclear antigen (LANA, ORF73), kaposin (K12), vFLIP (K13), vCyclin (ORF72), and 12 microRNAs (miRNAs) (3). Induction of viral replication and lytic gene expression, often by inflammation, promotes the expression of the Replication Transactivation Activator (RTA, ORF50), and a resulting cascade of secondary and tertiary viral proteins that make the virus capsid and DNA synthesis enzymes.

The etiological agent of Kaposi sarcoma (KS), KSHV exists in at least 5 subtypes (4) and is endemic in sub-Saharan Africa, parts of Eastern Europe, the Mediterranean, and parts of China, where rates can range from 30-90% (5). In the U.S. and many high-resource nations the prevalence of KSHV infection is low in the general population, but substantially elevated in high-risk groups such as HIV-1 infected individuals who have sex with men and in immunosuppressed subjects. Saliva is the major route of KSHV transmission and in endemic regions of the world, most infections occur within the first 5 years of life (6). KSHV is a class I carcinogen and about 1% of human tumors are associated with KSHV infection (7). Fewer than 1% of immunocompetent KSHV infected individuals develop disease, however most immunosuppressed individuals infected with the virus manifest one (or more) disorders (8) including KS, multicentric Castleman disease (MCD), KSHV inflammatory cytokine (KICS), immune reconstitution syndromes (IRIS), and primary effusion lymphoma (PEL).

KS is an incurable disease originally described as a blood vessel tumor in 1872 by Hungarian Moritz Kaposi, and is now understood to be a highly vascularized solid tumor of endothelial origin, characterized by KSHV-positive “spindle cells”, cellular pleomorphism, inflammatory infiltrate of lymphocytes and plasma cells, sinuous vascular spaces, extravasated erythrocytes, and fibrosis (9). Forms of KS include classical KS (cKS), iatrogenic immunosuppression-associated KS (iKS), endemic KS (enKS), and epidemic HIV-1/AIDS-associated KS (epKS) (10). The enKS and cKS are often indolent, whereas epKS can have widespread mucocutaneous, nodal, and visceral involvement (9). Although the majority of cells in a KS lesion manifest KSHV in latency, lytic reactivation is a critical step in oncogenesis (11). KSHV infection of endothelial cells or hematopoietic progenitors leads to changes in their morphology, glucose metabolism, proliferation, lifespan, and gene expression. KSHV oncogenicity is reflected by numerous pro-angiogenic molecules that are induced, including members of the vascular endothelial growth factor (VEGF)-VEGF receptor and angiopoietin families. Interleukin 6 (IL6) and IL8, and platelet-derived growth factor, through the activities of lytic proteins K1, K15, and viral G protein coupled receptor (vGPCR) (12). Latency proteins contribute to tumorigenesis through repression of apoptosis (LANA, vFLIP), and activation of cyclin-dependent kinase (vCyclin). KSHV also elaborates an array of mediators of immune evasion (13).

Although there are several transcriptomic studies of the latent and lytic KSHV genome, there is limited information from studies of KSHV-associated tumors (14). Given the cellular complexity, rarity of infected cells, and variable clinical course of KSHV associated disorders, single-cell analyses provide a unique opportunity to explore key interactions of lytic and latent infected tumor cells with the tumor microenvironment.

KS therapy is focused on disease palliation to improve quality of life and survival, but it is not curative (15). A key management component is to minimize immune suppression, whether by reducing immunosuppressive medications for iKS, or optimizing antiretroviral therapy for epKS. For indolent localized KS with minimal cosmetic or functional disturbance, topical or localized therapies may be indicated. For aggressive or visceral disease, or lesions with moderate-severe cosmetic or functional disturbance, systemic therapy is indicated. This may include FDA-approved chemotherapies such as liposomal doxorubicin or taxanes, or the immune or cereblon-modulator drug (IMiD, cel-mod) pomalidomide. The mechanism of action of these drugs remains unclear but they are known to alter angiogenesis, cytokine production, and T-cell activation (16). Other chemotherapeutic, anti-angiogenic, proteasome inhibitor, and immune checkpoint inhibitor drugs showed preliminary activity. However, response rates of 30-60% are seen with most approaches, and biomarkers of activity remain to be defined. In rare cases, exacerbation of KS-associated inflammatory disorders were seen (17).

Previous studies suggested that KSHV latent and lytic gene expression occurs in KS, and disruption of either program results in tumor regression (18). Animal models for KS are lacking, and there is a dearth of genomic studies on primary KS tissue due to the admixture of multiple cell types, the small proportion of KSHV positive cells, and the complexity of fibrotic skin tumors. Here we used a scRNAseq multi-omic approach to characterize the cellular and viral KS transcriptome at a single cell level in primary tissue.

These findings may have applications for discovery of prognostic and predictive biomarkers and therapeutic insights for the design of safe and effective therapies for KS. Application of these technologies to understand primary KS pathogenesis and therapeutic responses may be applied to understanding oncogenic virus biology, as well as defining the evaluation, and treatment of other infection-associated cancers.

## Results

Twelve participants contributed twenty samples for scRNAseq analysis, comprising fourteen KS skin biopsies, one non-KS skin biopsy, and five PBMC samples (Table 1). Eight HIV+ participants had AIDS-associated KS (epKS), one HIV-participant had classic KS (cKS), one HIV-participant had iatrogenic KS (iKS), one HIV+ participant who also had a renal transplant had KS (ep/iKS), and one HIV+ participant had neither KS nor AIDS. Three participants contributed samples before and after therapy, one treated with nivolumab and ipilimumab, one treated with antiretroviral therapy, and one treated with pomalidomide. The participant with classic KS was the only female in the cohort. The sample set is small and diverse and intended to demonstrate the feasibility and reproducibility of the methodology, generate hypotheses for future studies, and offer novel insights into potential biomarkers, pathways, and therapeutic targets.

**Table 1.** KS Primary Patient Samples.

| ID | Age | Sex | HIV Status | Subtype | CD4 count | Treatment |
| --- | --- | --- | --- | --- | --- | --- |
| KS1-PRE | 37 | M | POS | AIDS ASSOCIATED | 108 |  |
| KS1-POST | 37 | M | POS | AIDS ASSOCIATED | 156 | NIVO/IPI |
| KS2 | 26 | M | POS | AIDS ASSOCIATED | 139 |  |
| KS3 | 34 | M | POS | AIDS ASSOCIATED | <35 |  |
| KS4 | 32 | M | POS | AIDS ASSOCIATED | <35 |  |
| KS5 | 54 | M | POS | NOT KS | 444 |  |
| KS6A | 34 | M | POS | AIDS ASSOCIATED | 203 |  |
| KS6B | 34 | M | POS | AIDS ASSOCIATED | 347 | ART |
| KS7 | 81 | F | NEG | CLASSIC | >2000 |  |
| KS8 | 35 | M | POS | AIDS ASSOCIATED | 96 |  |
| KS9 | 31 | M | POS | AIDS ASSOCIATED | <35 |  |
| KS10-PRE | 61 | M | POS | AIDS ASSOCIATED | 225 |  |
| KS10-POST | 61 | M | POS | AIDS ASSOCIATED | 152 | POM |
| KS11 | 76 | M | POS | IATROGENIC | 114 |  |
| KS12 | 73 | M | NEG | IATROGENIC | N/A |  |

Cells utilized for scRNAseq were obtained from viably frozen single-cell suspensions prepared from fresh primary skin and blood samples and submitted in two batches for 10X Genomics 5’ gene expression with multiomic TCR profiling (Figure S1). KSHV transcripts could be detected in all KS skin tumors (and not in KS5, which is the KSHV-negative, dermal sclerosis sample). KSHV transcripts were not present in PBMC preparations. HIV-1 transcripts were not detected by this method in any sample. The rarity of viral reads in infected cells and the rarity of infected cells in tumors resulted in an analysis challenge in which stringent quality control measures typically applied to filter out noise (cells that expressed less than 100 features; genes that were expressed in less than 10 cells) also filtered out most KSHV-infected cells. Filters for dying cells (>20% mitochondrial genes) and low quality reads (proportion of UMI > 93^rd^ percentile) removed between 15 and 25% of cells from each skin sample (Figure S2). In skin samples, most noise resulted from 1 or 2 false-positive KSHV reads per cell and evaluating cells with more than 2 KSHV reads per specified gene retained 70% of suspected true positives and removed 99% of suspected false positives (Figure S3).

### The Landscape of Primary KS

To observe the cellular landscape of primary KS, 97,413 cells from eleven samples (4 PBMC samples and 7 skin samples) were merged into a single, dataset (Figure 1A, S2). Cells from the peripheral blood (n=38,266) were clearly distinct from those in the skin samples (n=59,147). In the peripheral blood, separable clusters of cells were found for monocytes and macrophages, neutrophils, B and T lymphocytes, and natural killer cells. Within skin biopsies, several clusters of endothelial cells and fibroblasts were detected as well as keratinocytes and epithelial cells. Clusters of T cells, B cells, macrophages, and dendritic cells were also detected in skin preparations and, interestingly, migrate more closely with clusters of the same cell types obtained from peripheral blood (Figure 1B). In addition to human transcripts, KHSV genes were also detected in 3,269 cells (5.5% of total skin tumor cells), exclusively from skin samples (Figure 1C), and clustered adjacent to but not within endothelial cells.

**Figure 1:**
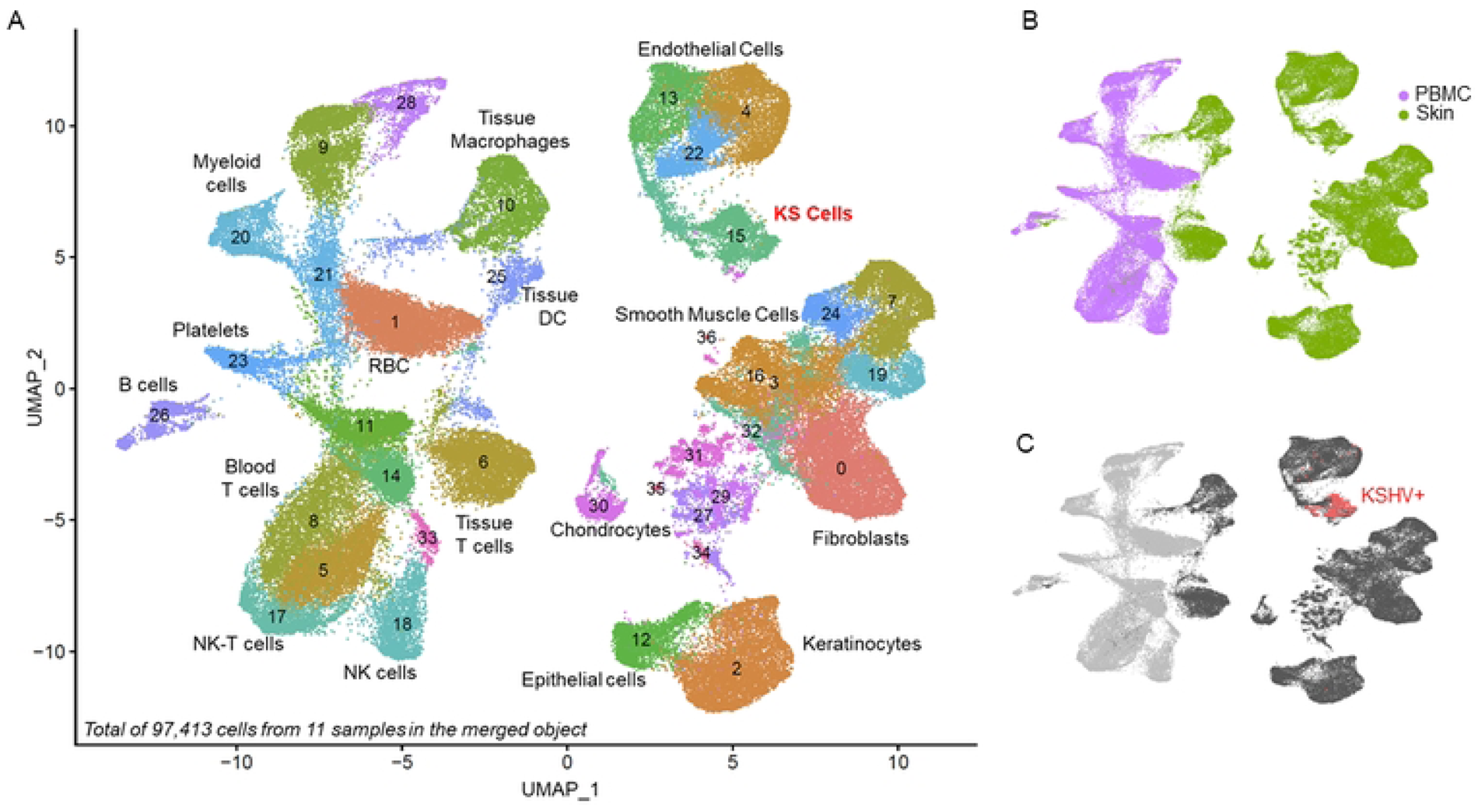
Landscape of primary KS. Single cell suspensions from viably frozen primary KS blood and tumor samples processed for scRNAseq resulted in, A) a merged UMAP plot of 97,413 cells from 11 samples representing a landscape of 36 clusters. Between 15 and 25% of cells from each sample were removed when filtering out dying cells (>20% mitochondrial genes), low quality cells (proportion of UMI > 93^rd^ percentile), and doublets (identified by DoubletFinder package). Cluster identities were annotated using the reference dataset from the human primary cell atlas. B) Cells obtained from skin (n=59,147) and PBMC (n=38,266) form clearly delineated clusters as well as tumor cells obtained before and after therapy from the same individual. C) Reads corresponding to KSHV genes can be detected in 3,269 tumor cells in the merged object corresponding to an average of 5.5% of total tumor cells. KSHV was not detected in PBMC samples.

### Two Populations of KSHV Infected Cells in Primary KS Skin Lesions

In KS skin tumors, cells in which KSHV genes were expressed formed two distinct clusters (Figures 2, S4). Both clusters contained cells expressing lytic and latent KSHV genes (Figures 3, S5) and both clusters express endothelial markers PECAM1(CD31), PDPN, LYVE-1, and CD36 (Figure S6). However, in each KS sample, one of the two KSHV-infected clusters contained cells with extremely high expression of both viral and cellular transcripts (Figure S7). Dozens to hundreds of host genes were also differentially expressed between these two infected cell clusters, including CD34 (Figures 4, S6, S7). A dramatic and unexpected distinguishing characteristic was the differential regulation of housekeeping genes including GAPDH and ACTB (Figure S8). In the CD34-cluster, housekeeping genes were suppressed while cellular proliferation factors were enriched including EP300 and CREBBP, along with factors like DTX1, DTX4, HEY1, and CTNNB1 that regulate the NOTCH and WNT signaling pathways (Figure 4). The KSHV gene vFLIP was elevated in this cluster of infected cells along with voltage-gated ion channels (Figure S8). Expressing biomarkers consistent with a lymphatic endothelial lineage, the CD34-cluster was likely representative of proliferating KS cells.

**Figure 2:**
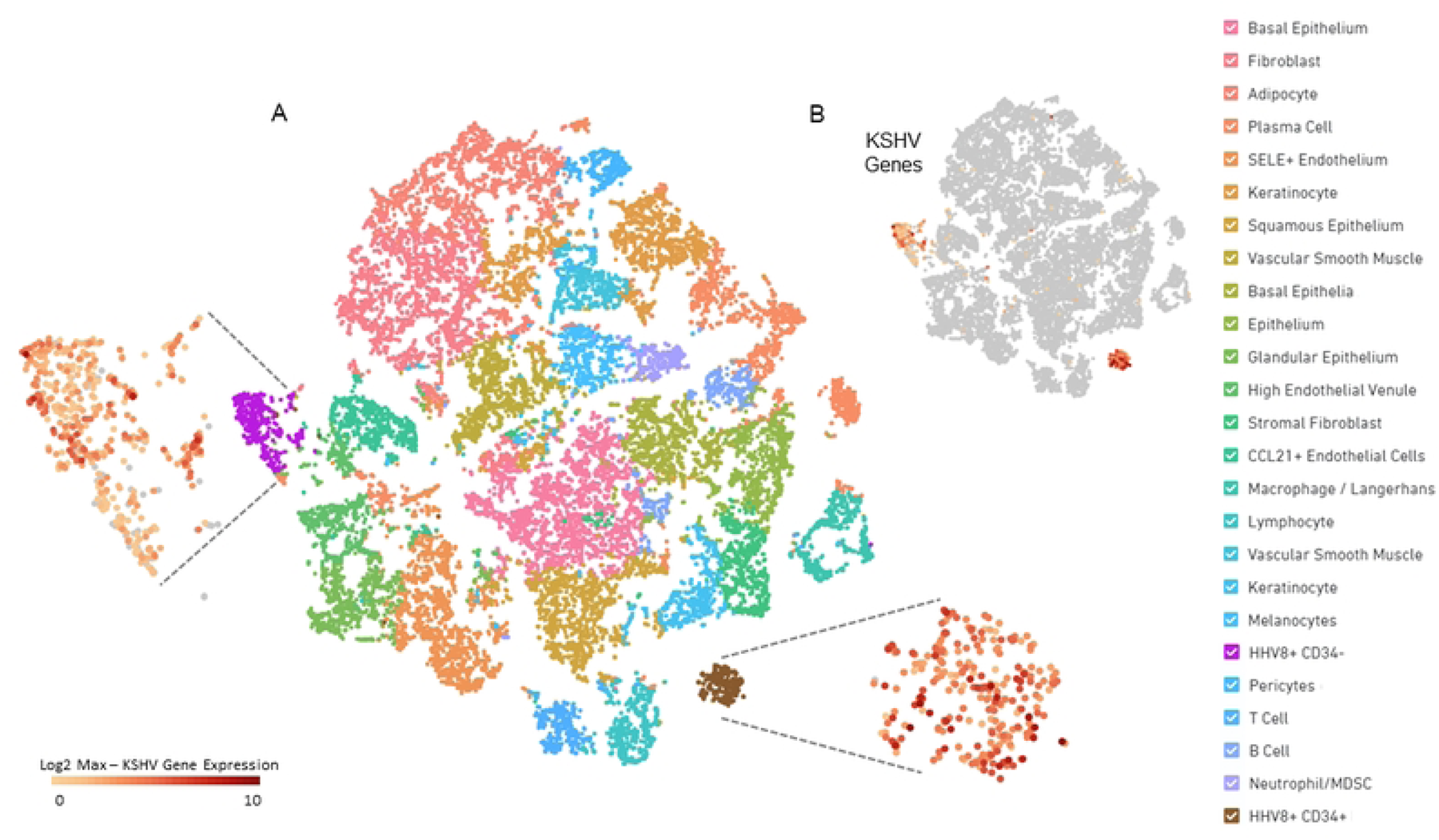
The Tumor Microenvironment of a Primary KS Skin Lesion. 10X Cell Ranger, A) graph-based t-SNE cluster plot of KS6B skin tumor with suspected cell type identities of clusters indicated in key. KSHV-infected cell clusters are highlighted (purple, brown) alongside expanded insets color coded to show Log2 KSHV gene expression in each cell of the cluster or B) KSHV gene expression in the entire sample.

**Figure 3:**
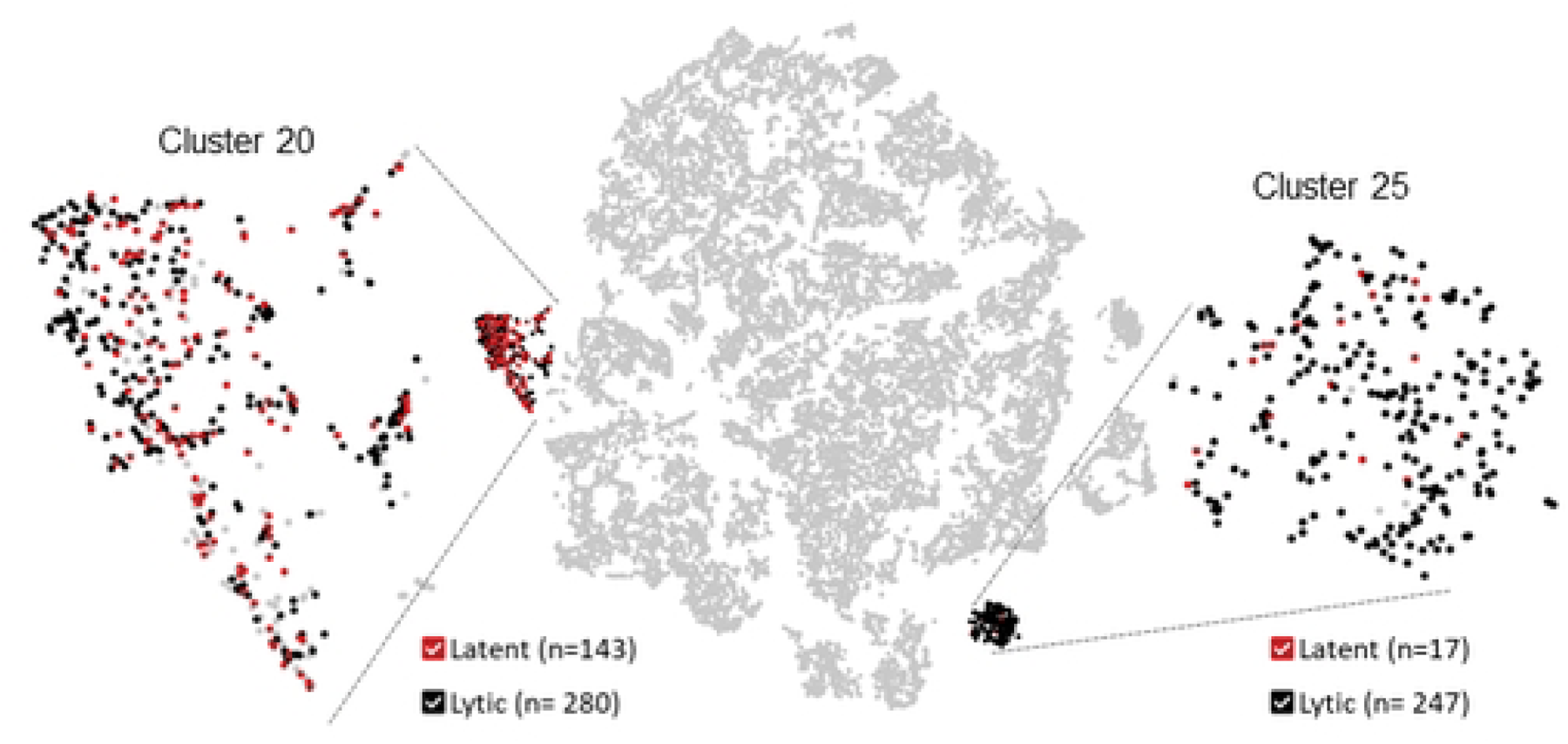
Detection of Lytic and Latent Cells in Primary KS lesions. Graph-based t-SNE cluster plot of KS6B skin tumor revealing tumor cells carrying latent KSHV. Latency is defined as cells expressing any of 6 genes expressed in latency, LANA (ORF73, gp81), Kaposin (K12, gp79), vFLIP (ORF71, gp80), K15 (ORF75, gp85), vOX-2 (K14, gp83), vIRF-2 (gp65) and not expressing any other KSHV genes. Cells engaged in KSHV+ lytic replication is defined as cells expressing any KSHV genes but excluding cells defined as Latent.

**Figure 4:**
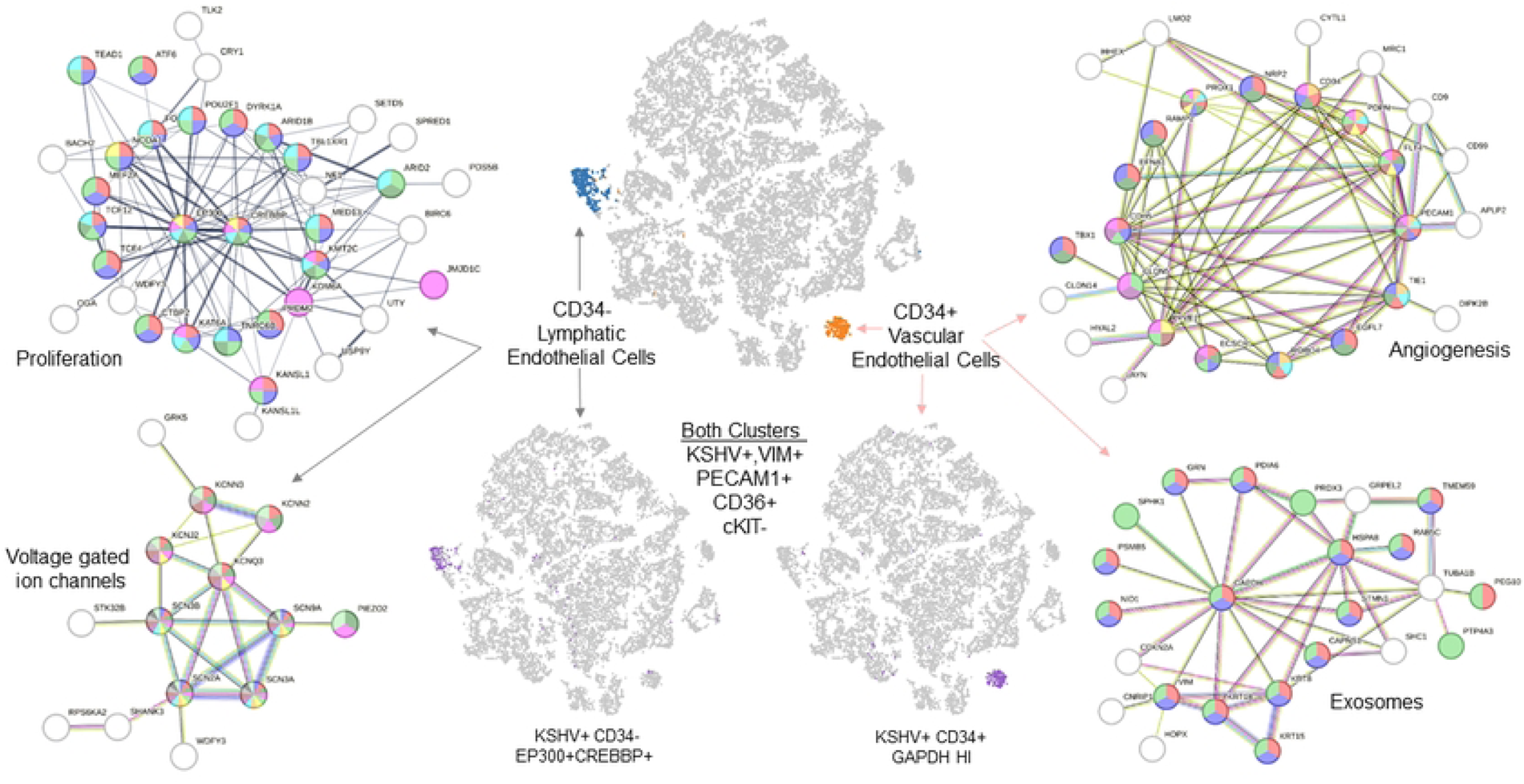
Two Populations of KSHV Infected Cells in Primary KS lesions. Differential gene expression profiles for two KSHV infected cell clusters revealed similarities between the two t-SNE clusters including the presence of KSHV genes as well as VIM, PECAM1, and CD36, and the absence of cKIT in both clusters. However the KSHV+ clusters diverged dramatically into CD34-lymphatic endothelial cells (black cluster) and CD34+ vascular endothelial cells (red cluster). The CD34-population was enriched in proliferation factors including EP300 and CREBBP, genes associated with NOTCH and WNT signaling, voltage-gated ion channels (shown as STRING objects), and low expression of housekeeping genes including GAPDH, possibly due to virus host shut off. The KSHV gene vFLIP was also elevated in the CD34-cluster. The CD34+ cluster was enriched in CD90 and S100A6 and genes related to angiogenesis and production of extracellular vesicles. Unlike the CD34-cluster, these cells expressed very high levels of housekeeping genes including GAPDH as well as factors associated with ribosome biogenesis and electron transport and the KSHV gene K5. The CD34+ cluster of KSHV infected cells may be KS spindle cells.

The CD34+ cluster of KSHV expressing cells exhibited high expression of GAPDH and ACTB as well as CD90, PROX1, CD36, PDPN, LYVE1, and the viral gene K5 (Figure S5-S8). CD34 expression in this cluster in the setting of endothelial marker expression is consistent with blood vascular endothelial cell identity. (Figure 4).CD34+ vascular endothelial cells can be distinguished from CD34+ telocytes by the expression of CD31+, PDPN+, and LYVE+, and the absence of PDGFRA (19). The CD34+ KSHV+ cluster expressed transcripts associated with ribosome, spliceosome, and electron transport machinery and also exhibited high expression of lymphocyte antigen 6 complex, locus H (LY6H) which has been described by Moorad et al. as associated with “inflammatory” KS lesions (Fig S9) (20). Despite the significant differences in viral, cellular, and specific biomarker gene expression, the two infected clusters were grouped together in our merged UMAP plot combining all 11 samples’ data, supporting similarities between these 2 clusters that could be due to a common endothelial cell lineage origin. Interestingly, both clusters of KSHV-infected cells were present in KS tumors from all participants, both may be involved in tumor growth, and each cluster represents a largely uncharacterized and rare sub-population of cells within KS tumors that are readily distinguishable by expression of housekeeping genes.

### Novel Biomarkers of KSHV infected cells in primary KS lesions

In addition to viral genes, several cellular genes were commonly expressed in the KSHV-positive cells including prospero homeobox protein 1 (PROX1), mannose receptor C-type 1 (MRC1; CD206), fms-related tyrosine kinase 4 (FLT4), and Kir2.1 inward-rectifier potassium channel (KCNJ2) (Table S1). These markers have been previously described in KS lesions and provide confirmation for the sensitivity and reproducibility of scRNAseq data obtained from primary skin lesions (21–23). Differential expression analysis revealed 1,022 significantly enriched genes in KSHV+ cells in Cluster 15 cells including several voltage gated ion channel (VGSC) genes with SCN9A as a top biomarker candidate along with KSHV genes LANA and Kaposin (Figure 5, Table S1). The expression of the VGSC gene SCN9A has not been previously described in KS, was tightly associated with both clusters of KSHV infected cells (Figures S10, S11), was minimally expressed in uninfected endothelial or other stromal cells, and was not expressed in the KS-negative skin sample.

**Figure 5:**
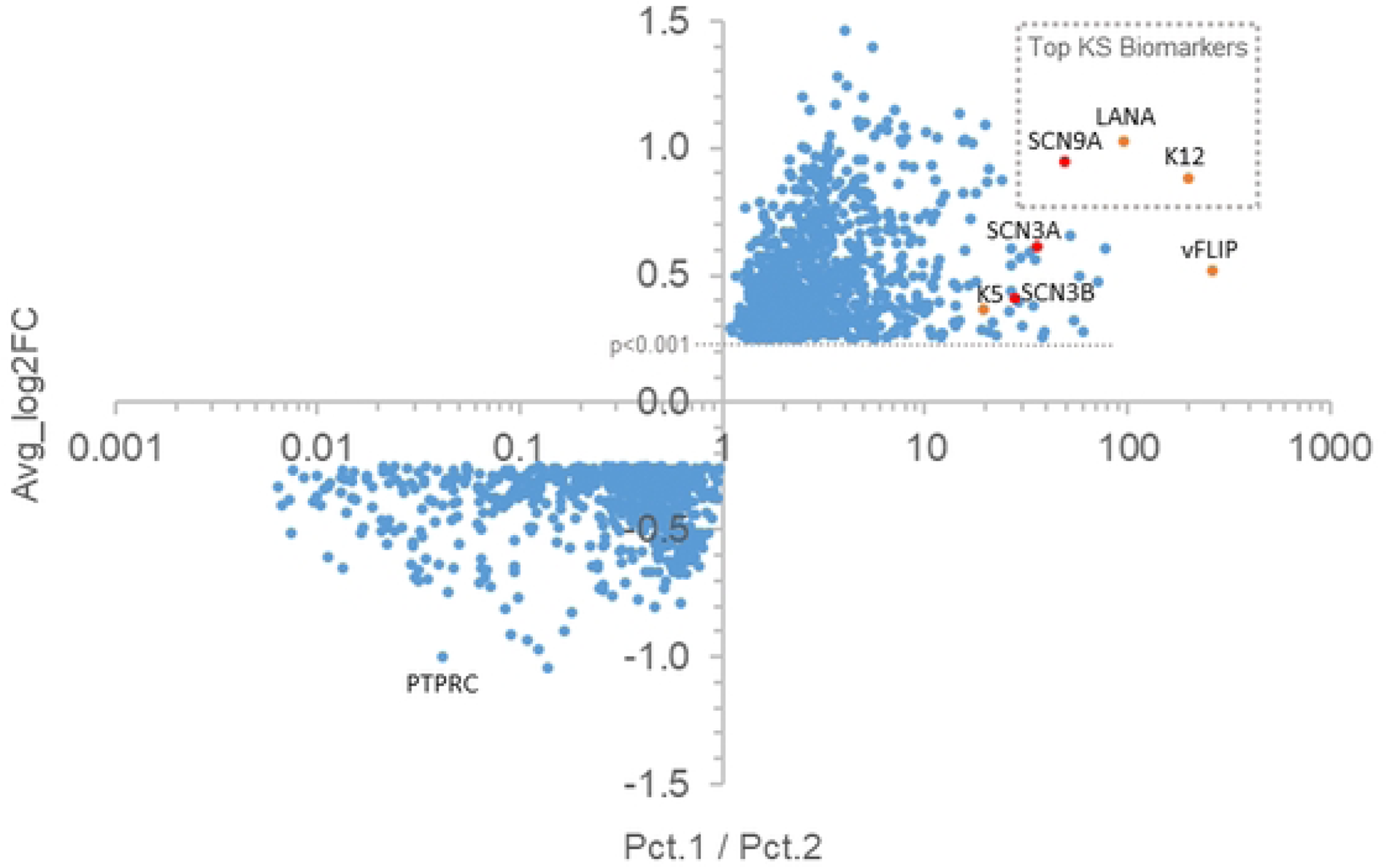
Differential Expression Analysis of KSHV+ cells. Differential expression analysis of genes expressed in KSHV+ cells within UMAP cluster 15 (Figure 1B; average Log2 fold change vs KSHV negative cells in all other clusters) plotted against the ratio of percent of KSHV+ cluster 15 cells expressing that gene (Pct.1) to the percent of KSHV-cells expressing that gene in all other clusters (Pct. 2). The KSHV genes LANA and K12 were the top viral biomarkers and the voltage-gated sodium channel SCN9A (Nav1.7) emerged as the top, non KSHV biomarker.

### The number of KSHV+ cells is inversely proportional to immune cell number in primary KS

In several samples, including two HIV-negative samples, KSHV+ cells were very rare, 1% or less of the total cells in the sample (Figure 6A). In other samples, all of which were HIV-associated, KSHV+ cells were significantly more abundant, representing 3-7% of the total cells in the sample. In samples in which KSHV+ cells were rare, macrophages expressing IL10 and IL1B, and CD8+ T lymphocytes were significantly more abundant than in samples in which KSHV+ cells were prevalent (Figure 6B). In these samples there was a strong inverse correlation between the number of IL1B expressing cells and the number of KSHV+ cells in the primary tumor (Figure 6C) suggesting a critical role for skin resident IL1B expressing cells in KS immunity.

**Figure 6:**
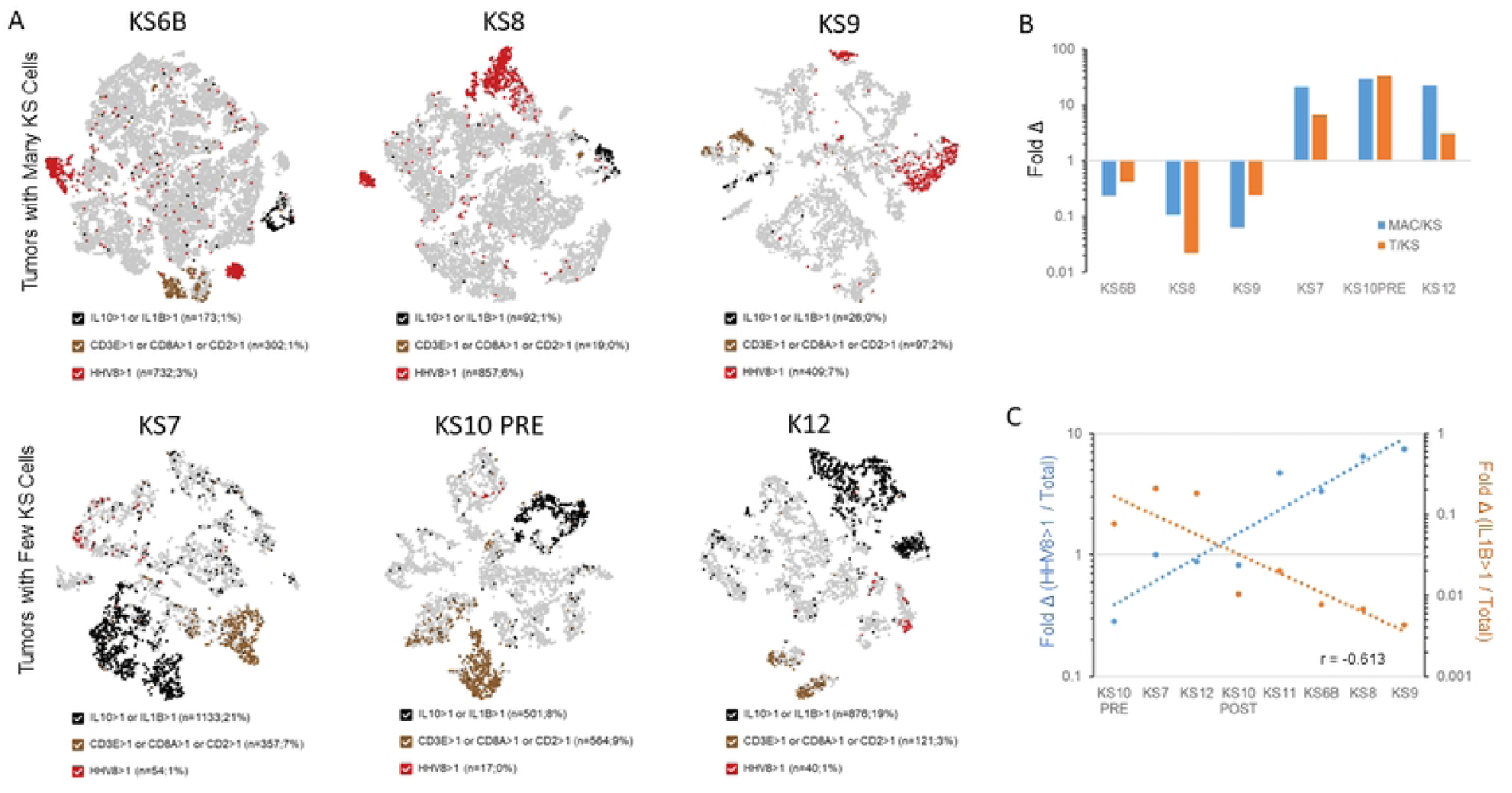
KSHV+ cell number is inversely proportional to immune cell number in KS skin tumors. A) t-SNE plots of primary KS tumor samples highlighting Macrophages (Black dots; IL10, IL-1B), T cells (Brown dots; CD2, CD3E, CD8A), and KSHV+ tumor cells (Red dots, KSHV genes). The upper three samples have more KSHV+ cells (3.4%-7.4%); the lower three samples have fewer KSHV+ cells (0.3%-1.0%). When poor quality and dying cells and doublets are removed from the data sets the ranges of KSHV+ cells in each group become 7.5-12.1% and 0.5-1.75%, respectively. B) Graph representing the ratio of Macrophages to KSHV+ cells (blue bars), and T cells to KSHV positive cells (orange bars) in primary KS skin tumors revealing that in tumors in which KS cells are rare, Macrophages and T cells are significantly more abundant. C) Double Y-axis graph (Left axis / Blue line: Ratio of KSHV+ cells to Total cells; Right Axis / Orange line: Ratio of IL1β+ cells to Total cells) demonstrating the inverse correlation between IL-1β and KSHV.

### The Peripheral Blood of KS Subjects

Whether truly absent or simply below the level of detection for scRNAseq, there were no detectable HIV-1+ or KSHV+ reads in the peripheral blood mononuclear cells (PBMCs). However, the data did reveal unique characteristics of T cells in the peripheral blood of KS patients.

### Low CD4:CD8 ratio in Peripheral Blood in HIV+ KS Subjects

The CD4:CD8 T cell ratio is an important biomarker of pathogenesis. In healthy subjects the CD4:CD8 T cell ratio is usually greater than 1.0, indicating that CD4+ T cells are typically present in greater abundance than CD8 cells (24). In the peripheral blood of KS patients, the CD4:CD8 T cell ratio is less than 1, indicating either a loss of CD4 cells or a gain of CD8 cells or, more dramatically, both (Figure 7). This is a hallmark of AIDS, resulting from persistent HIV infection and consistent with the characterization of KS as an AIDS-defining illness. Accordingly, the KS12 blood sample from the HIV-negative iKS participant, had the highest CD4:CD8 T cell ratio, albeit still <1, largely due to the dearth of both CD4+T cells and CD8+T cells. In calculating these ratios, the value of combining scRNAseq with TCR sequencing was apparent, clearly distinguishing TCR-CD4+ monocytes from TCR+CD4+ T cells, and TCR-CD8+ NK cells from TCR+CD8+ T cells.

**Figure 7.**
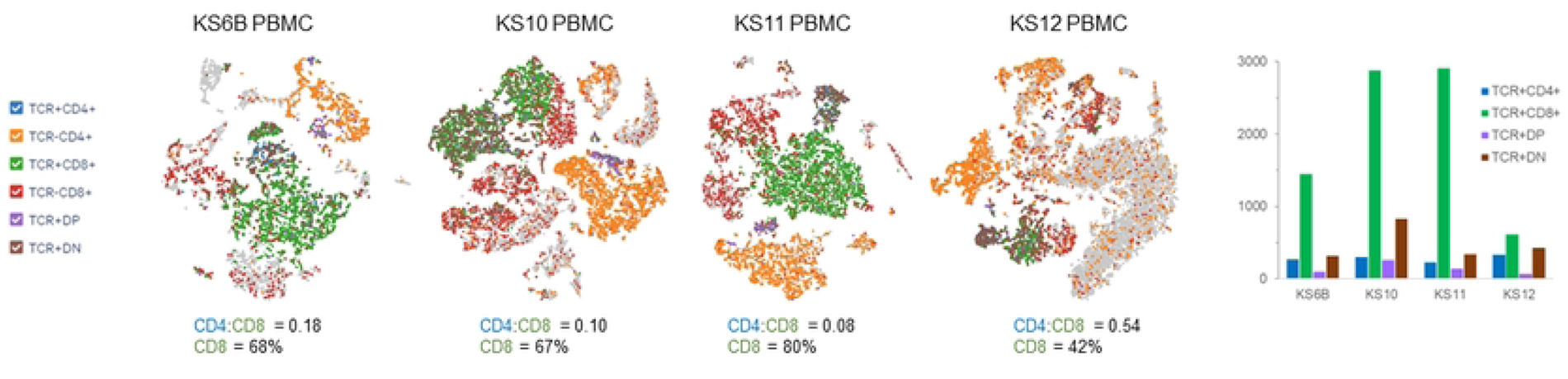
The CD4:CD8 Ratio is very low in the peripheral blood of KS patients. t-SNE plots for 4 primary PBMC samples showing clusters representing the following cell types: TCR+CD4+ (Blue; CD4+ T cells), TCR-CD4+ (Orange; monocytes), TCR+CD8-(Green; CD8+ T cells), TCR-CD8+ (Red; NK), TCR+DP (Purple; double-positive T cells), TCR+DN (Brown; double-negative T cells). The ratio of CD4+T cells to CD8+ T cells is indicated, as well as the percentage of CD8+ T cells out of total TCR+ cells

### Expansion of CD8 T cell clones in KS Subjects

T cells present in the peripheral blood exist as a unique oligoclonal clonal pool of various T cell clones carrying a diverse array of antigen-specific T cell receptors (TCRs) (25). An emerging challenge in tumor immunology is elucidating the role and identity of tumor or pathogen specific CD8 T cell clones. By combining single-cell TCR profiling with single-cell gene expression, multiplex scRNAseq provides a powerful tool to identify KS-specific CD8+ TCR clones (26). In eight samples in which TCR reads from matched tumor and PBMC from four participants could be evaluated, the distribution of VDJ recombination was non-random among the most abundant CD8+ clones (Figure 8). Identical T cell clones were found in both the skin and the peripheral blood from the same patient and in two different skin samples taken before and after therapy from the same patient (Figure 8A, S12). Interestingly, in these cases, the most abundant T cell clone in the peripheral blood was not the most abundant clone in the tumor. The most abundant T cell clones among the KS samples carried a subset of frequent rearrangements, especially enriched in TRBJ1-1 and TRBJ2-5, with recurring similarities among the CDR3 sequences of the most abundant clones in each patient. Interestingly, TRBJ1-1 and TRBJ2-5 were not the most frequently used TRBJ variants in the TCRβ repertoire of healthy controls described by Drulak et al. (27), and the CDR3 sequences associated with TRBJ1-1 clones shared significant similarities with the CDR3 sequence (CASSILGLRNTEAFF) found in CD8+ T cell clones that react with the KSHV major capsid protein (ORF25) (Figure 8C) (28). These data are novel, and they suggest the importance and therapeutic potential of KSHV-targeted CD8 T cells.

**Figure 8.**
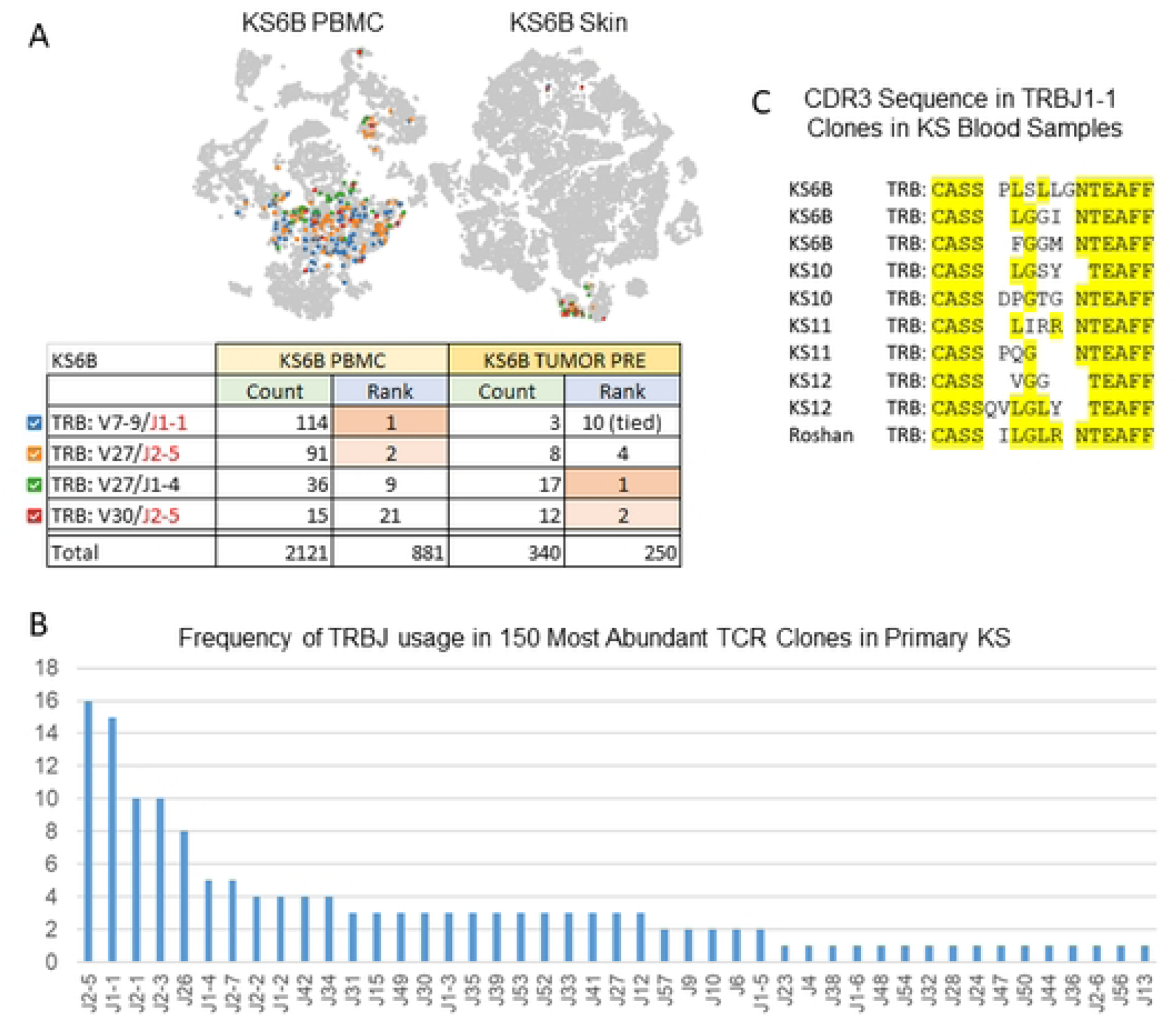
Expansion of CD8 T cell clones in primary KS. A) t-SNE plots revealing the two most predominant TCR clones in the peripheral blood (Blue and Orange) and skin tumor (Green and Red) from the same patient (KS6B). B) Graph depicting the frequency of TRBJ usage in the 150 most abundant TCR clones in primary KS samples with matched blood and skin samples (KS6B, KS10, KS11, KS12). C) CDR3 sequence in TRBJ1-1 clones in KS PBMC samples compared to the sequence reported in Roshan et al.

### Tumor-Associated KSHV Single Cell Transcriptome

#### Detection of Lytic and Latent KSHV gene expression in primary KS skin lesions

In primary KS lesions, only a small subset of cells were positive for KSHV. KSHV gene expression is tightly regulated in infected cells and the virus persists in lytic and latent phases of replication and dormancy (3). During latency, the virus is largely quiescent, and expresses a small subset of viral genes including LANA (ORF73; gp81), Kaposin (K12; gp79), vFLIP (ORF71, gp80), K15 (ORF75, gr85), vOX-2 (K14, gp83), and vIRF-2 (gp65). Cells in which any or all of these genes were expressed, but not other KSHV genes, were considered to be harboring latent virus. Cells in which latency genes together with other KSHV genes were expressed were considered to be harboring virus in lytic replication. LANA was generally the most highly expressed latency gene, and K5 was the most highly expressed lytic gene (Figures S5, S8). One sample, KS6B, may have an amplification of the portion of the viral genome encoding K5-K7 (Figure S13). Amplification of this region of the KSHV genome has been described in approximately one-third of virus samples harvested from primary KS tissues (29).

#### Detection of KS-specific Viral Transcripts and Quantitation of Viral Load

Utilization of single cell RNAseq, bulk RNAseq, and DNA-based quantitation assays on a single primary sample not only compensates for limitations of each methodology but also offers complementary insights and quality control for each sample. For example, Kaposin mRNA is detectable in scRNAseq data, but splice variants of Kaposin transcripts have been described in KS that may be difficult to detect by scRNAseq, or quantitate via ddPCR. We utilized probe-capture RNAseq for detection and quantitation of viral genes involved in lytic and latent viral gene expression in bulk RNA from a primary lesion and also for the detection of specific splice variants of Kaposin mRNA (Figure S14A). Similarly, in order to rapidly quantify KSHV viral load (the relative abundance of viral genomes per cell in a sample), primers and a probe were designed for a KSHV-specific digital droplet PCR assay (Figure S14B). These tools were utilized to quantitate viral DNA and RNA in primary samples in addition to scRNAseq studies.

#### Evaluation of Therapeutic Interventions

By evaluating samples obtained before and after therapeutic intervention, scRNAseq can be used to identify the effects on viral gene expression, tumor cell abundance and gene expression. In addition, scRNAseq can assess the impact of therapy on the abundance and expression of cells in tumor stroma and peripheral blood. As a proof of principle, we evaluated one pair of serial samples, each before and after introduction of antiretroviral therapy (Figure 9), nivolumab and ipilumumab therapy (Figure S15), or pomalidomide therapy (Figure S16). In one HIV+ KS participant, the tumor sample obtained after 8 months of antiretroviral therapy had significantly more CD8 T cells and increased stromal expression of VEGFC (FLT4 Ligand) and decreased expression of IL-6 (Figure 9). In a second HIV+ participant, the abundance of activated CD8+ T cells in the tumor harvested after nivolumab and ipilumumab therapy was greater than in the skin tumor harvested prior to therapy (Figure S15). Moreover, the number of cells expressing Kaposin, but not those expressing vFLIP, were decreased. In a third HIV+ participant, in the tumor sample harvested after pomalidomide therapy there were significantly more KSHV-infected cells expressing high levels of LANA than in the tumor harvested prior to therapy (Figure S16). Interestingly, there were also fewer activated T cells in the tumor harvested after therapy.

**Figure 9:**
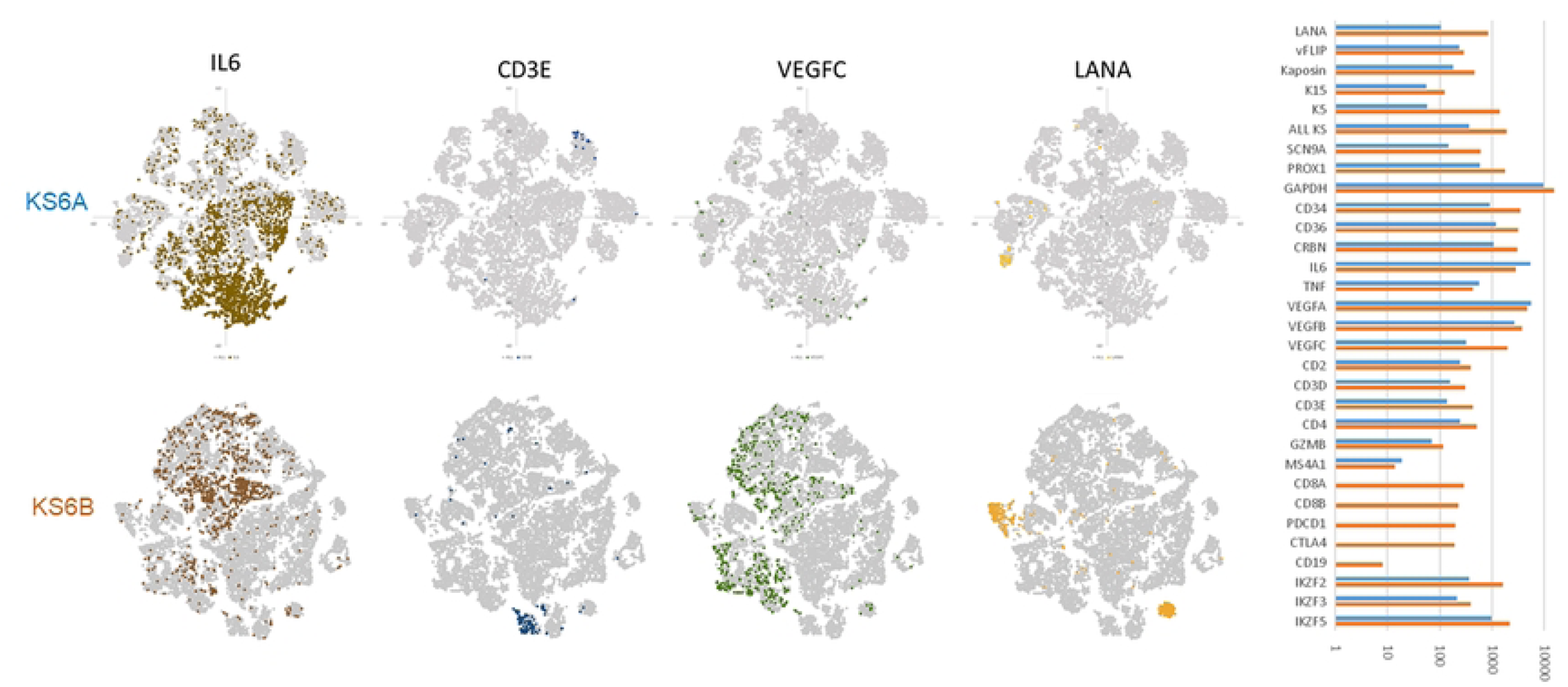
Evaluation of Serial Samples Reveals Changing Tumor Environment. KS biopsy samples obtained from the same patient (KS6), 9 months apart, after CD4 count in peripheral blood had rebounded to 347 in response to antiviral therapy. Cells positive for IL6, CD3E, VEGFC, and the KSHV gene LANA are shown in t-SNE plots before (KS6A) and after therapy (KS6B). Read counts for selected genes before (blue bars) and after therapy (orange bars) are shown in the graph on a log scale.

## Discussion

KSHV is an ancient human oncovirus and KSHV-associated diseases, such as KS, PEL, and MCD exhibit widely different transcription programs (30–32). Bulk RNAseq studies of transcription start sites were found to be cell-type specific in start site usage and promoter strength (33). Lidenge et al found similar profiles in enKS and epKS, although genes involved in tumorigenesis and inflammatory and immune responses were more highly expressed in enKS (34). They also showed that antiretroviral use and gender had little impact on the KS transcriptome. Dittmer *et al* examined epKS biopsy specimens and found KSHV lytic mRNAs in only 1 of 8 samples from HIV-suppressed individuals, compared to 7 of 11 of biopsies from subjects with fulminant AIDS (31), noting high levels of expression of viral genes K1, viral G protein coupled receptor (vGPCR, ORF74), and vIRF1. RNAseq studies of Tso *et al*. on four epKS skin biopsies from subSaharan Africa showed high levels of expression of viral immune modulation genes, vIL6 (K2), modulator of immune recognition (K5), viral inhibitor of apoptosis (K7), and ORF75 (35). Robust lytic gene expression was found in 2 of 4 KS tumors. They also noted upregulation of transforming growth factor-beta 1 (TGFB1), and chemokine receptor CXCR3 and ligands CXCL-9, -10, and -11. Infiltration of B lymphocytes, macrophages, and NK cells was found in all samples, but dendritic cells in only 2 cases. Activation of glucose metabolism genes was coupled with decreased expression of lipid anabolic and catabolic genes. Gjyshi *et al*. found overexpression of the nuclear respiratory factor 2 (Nrf2) associated with repression of the latent-lytic switch in infected PEL cell lines (36). Rose *et al.* published a comprehensive sequence analysis of 41 epKS tumors from 30 individuals in Uganda, all naïve to antiretroviral therapy (30). This study revealed three clusters of tumors with different latent and lytic KSHV gene expression profiles. They noted that tumors with a latent phenotype had high levels of total KSHV transcription, while tumors with a lytic phenotype had low levels of total KSHV transcription. They noted no difference in transcription profiles of morphologically distinct tumors from the same individual. In addition, they found no correlation between levels of KSHV transcripts/cell and the number of copies of KSHV genomes/cell.

Several recent studies have applied scRNAseq to acute virus infections, such as HIV, influenza virus, or flaviviruses (37–41). scRNAseq has also been used to analyze latent infection with human cytomegalovirus and other herpesviruses (42–44). In a previous scRNAseq study of KSHV infected PEL cell lines, Landis *et al*. found latency-associated transcripts in the majority of cells of two PEL cell lines (45). Fewer than 1% of cells expressed lytic viral RNAs, with a predominance of early over late lytic viral transcripts. Jung *et al.* performed scRNAseq on KSHV infected 3-dimensional air-liquid interface organoid cultures, which permitted high levels of lytic replication, and a unique pattern of lytic K2-K5 gene expression, accompanying marked changes in host gene expression in infected and uninfected cells in different epithelial layers (46).

The current study provides the first single cell transcriptomic analysis of primary KS tumors. Our study included samples from different subtypes of KS, including cKS, iKS, and epKS. In addition, we included several peripheral blood mononuclear cell preparations obtained at the same time as skin biopsies. The single cell transcriptomic profiles defined distinct clusters of cells in the blood and tumors, including hematopoietic cells, and tumor-associated fibroblasts, vascular smooth muscle cells, melanocytes, keratinocytes, and several distinct populations of epithelial and endothelial cells.

In this study, KSHV transcripts were found in a minority of cells in each KS tumor but not in cells in the peripheral blood. PCR assays have detected KSHV DNA in peripheral blood mononuclear cells in 52% of epKS subjects (47) and in studies of cKS, eKS, iKS, and epKS, detection of KSHV DNA in peripheral blood was associated with KS risk (48–51). Our failure to detect KSHV transcripts in peripheral blood mononuclear cells could be due to low levels of expression, a limitation of the sensitivity of our current single cell RNA sequencing technology, or both. Future studies using CITE-Seq with a custom panel of KSHV antibodies could improve sensitivity of this analysis.

A novel finding from our study was the consistent presence of two separate clusters of KSHV-infected cells in tumors, differentiated by the presence or absence of CD34 expression. The CD34-cluster was consistent with lymphatic endothelial lineage and was characterized by high levels of proliferative gene expression and voltage-gated ion channels. The CD34+ cluster of KSHV-infected cells are likely vascular endothelial cells and expresses many genes that correlate specifically with endothelial “tip” cells which drive angiogenic sprouting, are motile, and express long filopodia (52). Cells in these clusters shared common biomarkers of endothelial cells, including PECAM1 and CD36. However, in addition to marked differences in viral and cellular gene expression, we observed differential expression of several biomarkers that distinguish BVECs from lymphatic endothelial cells (LECs), such as CD34. We conjecture that challenges in developing KS tissue culture models may be due to lack of cultivation conditions that include both the CD34-and CD34+ KSHV infected cell types.

Interestingly, a study proposed a mechanism of “transcriptional reprogramming” in which PROX1 overexpression leads to suppression of BVEC gene expression, and induced the LEC transcriptional program (53). As discussed above, PROX1 is a known upregulated gene in KS, and we found consistent upregulation of PROX1 in both clusters of KSHV-infected cells. This finding reinforces the potential lineage relationship between the two infected clusters observed in our study, and it suggests PROX1 as a potential mediator of cell differentiation and/or malignant transformation in KSHV pathogenesis.

Although latency-associated transcripts predominated, lytic transcripts were also identified in both KSHV-infected cell clusters in our study. A critical role of lytic gene expression in KS was first proposed by Ganem (18). Although we cannot exclude some level of KSHV reactivation during processing of tumor samples, we noted the consistent finding of lytic transcripts in all cases. The most abundantly expressed viral genes in our samples were LANA, a latency-associated episome persistence gene, Kaposin which is a pathogenesis factor that encodes at least three proteins, one of which can drive host cell proliferation, and K5 which is an early lytic transmembrane ubiquitin ligase with immune evasion functions (54). Three other latent program genes were amongst the most expressed viral genes in our sample (gp80, gp83, vIRF-2), with CD34-clusters showing higher average reads per cell than CD34+ clusters. One notable exception was K5, which was slightly more highly expressed in CD34+ clusters, while being predominant in both clusters. A potential explanation for this early lytic gene being overrepresented in all clusters could be related to genomic rearrangements, which were not fully investigated in this study. One study showed K5 overexpression in almost a third of KS lesions, due to several *de novo* mutations resulting in similar KSHV genomic rearrangements of a 1.5kb section containing the K5 and K6 genes (29).

Differential expression of host genes, such as GAPDH and CD34, in KSHV infected cell clusters did not correlate with the latent or lytic viral programs in our study. Pardamean et al described two lytic-associated mechanisms of KSHV host gene shutoff, involving vSOX and ORF10 expression during early and late lytic replication, respectively (55). We detected no vSOX or ORF10 transcription in any of our samples, and we found notable reductions in GAPDH and other host gene expression in infected clusters with low lytic program gene expression. It is unclear whether host gene shutoff is induced by vSOX and/or ORF10 expression below the level of detection for our assay, or other host gene inhibition mechanisms are at play.

We also found that the number of KSHV+ cells in tumors was inversely proportional to immune cell number. It is notable that Landis et al noted different subpopulations of KSHV transcripts, even within a single latently infected PEL cell line (45). They found that the majority of cells only expressed canonical viral latent transcripts, a minority of cells exhibited more permissive transcription, and in some cells, no KSHV transcripts were detected. It should be noted that our study design would not have identified the latter population of cells with KSHV DNA, but no viral transcripts. Dittmer reported that in HIV-suppressed patients on antiretroviral therapy, KS lesions exhibited almost exclusively latency-associated transcripts, whereas the more permissive transcription pattern was identified in early AIDS KS lesions (31, 32).

Our study identified several potential cellular biomarkers of KSHV infection, including some that were previously described (21–23). One of these markers, PROX1, co-localizes with CD34 in KS lesions (21) and in other cancers (56), and may be involved in regulation of endothelial to mesenchymal transition. Another marker, FLT4, has also been linked to malignancy (57) and discussed as a potential therapeutic target (58).

The expression of VGSCs in both clusters of KSHV-infected cells was a novel observation. VGSCs consist of a main α subunit forming the channel, associated with one or two β subunits (59). VGSCs are abnormally expressed in many types of cancers, and their level of expression and activity are related to the aggressiveness of the disease. Their effects on tumor cell migration and invasiveness may be mediated through effects of sodium ions, through modulation of membrane potential, or other pathways (60). Of particular note was SCN9A (Nav1.7), a tetrodotoxin-sensitive VGSC with known roles in angiogenesis and regulation of chemotaxis (61). SCN9A is normally expressed in skin vasculature (62), as well as dorsal root ganglion cells and peripheral neurons (63). SCN9A has also been associated with several types of cancers, including endometrial, gastric, and prostate carcinomas (64–66). In this study, SCN9A was tightly correlated with viral gene expression and is a top biomarker candidate of KSHV infection within primary KS lesions.

A recent study by Dittmer et al identified two types of KS lesions, inflammatory and proliferative, based on host gene transcription patterns in bulk RNA sequencing (20). Of the transcripts associated with the inflammatory subtype, we detected cell migration-inducing and hyaluronan-binding protein (KIAA1199; CEMIP) expression in all KS tumor samples, in both CD34+ and CD34-clusters. CEMIP was among the most commonly expressed genes in the KSHV infected cells. Vesicle amine transport protein 1 homolog (T. californica)-like (VATL1) was also expressed in both infected cluster subtypes, to a lesser level and not in all samples; and LY6H was enriched in CD34+ clusters. Interleukin 1β (IL1B) was expressed in non-infected macrophages. Additionally, the proliferative subtype transcript annexin A8-like 1 (ANXA8L1) was detected in non-infected epithelial cells and keratinocytes in all of our samples. These single cell data characterize the differences in the transcription landscape within primary KS lesions and provide further insight into the potential biomarker function of these transcripts.

Among the KSHV-negative cell population in KS tumors were T cells. CD8 T cells are major mediators of anti-viral immunity and can rapidly recognize and kill cells expressing KSHV lytic antigens (67). KSHV-specific CD8 T cells are more frequent in KSHV-seropositive individuals without KS than those with active disease, suggesting an anti-tumor role of these cells (68). In the current study, we assessed the TCR repertoire of CD8+ T cells in KS tumors and blood samples and, in distinction to previous reports, identified clonally expanded T cells in tumors (69, 70). Clonally expanded CD8+ T cells could be detected in paired samples of tumor and blood as well as longitudinal samples of tumor. TCR Vβ CDR3 sequences of CD8 clones in the peripheral blood of KS were similar, but not identical, to a clone known to be directed against the major KSHV capsid protein (28). Differences in the abundance of CD8+ clones in blood and tumor may represent differences in affinity, avidity, invasiveness, or survival of disparate T cell populations, which could be directed against KSHV, cellular tumor-specific antigens, HIV-1, or other persistent viral infections in these individuals.

A comprehensive understanding of how transcriptomic alterations correlate with the microenvironment, cell type, disease type and severity, and response to therapy could yield predictive biomarkers. Tumors with infiltration of pro-inflammatory immune cells and CD8+ T cells may confer a better prognosis, whereas those with an abundance of regulatory T lymphocytes (Tregs) or myeloid-derived suppressor cells (MDSCs) often correlate with worse outcomes (71). The communication between tumor cells and the tumor microenvironment (TME) that regulates the dynamic balance between immunotolerance or immune rejection is the ultimate target of immunotherapy. A study of hepatocellular carcinoma identified a novel CD8+ T cell signature governed by layilin expression that led to exhaustion through inhibition of interferon (IFN)-γ production (72). In breast cancer, scRNAseq was used to identify a new class of tissue-resident memory CD8+CD103+ T cells with pro-inflammatory and cytotoxic characteristics (73). A lung cancer scRNAseq study identified a highly migratory T cell cluster linked to a positive response to immune checkpoint inhibitor therapy (ICT) (74), and another study identified Myc expression in endothelial cells as a contributing factor to tumor angiogenesis (75). Similarly, scRNAseq analysis of human papilloma virus (HPV)-associated carcinomas highlighted the role of B cells in ICT responses (76).

As a proof of the potential feasibility of scRNAseq to monitor KS therapy, we examined samples obtained longitudinally from three individuals. In one individual each, we assessed the effects of antiretroviral therapy, immune checkpoint inhibition, and pomalidomide therapy. With antiretroviral therapy and immune checkpoint inhibitor therapy, we noted a significant increase in tumor infiltrating CD8+ cells. In contrast, with pomalidomide therapy, we noted a decrease in tumor infiltrating activated T cells. Studies of additional individuals are warranted to comprehensively assess effects of immunotherapy on the KS tumor microenvironment.

One limitation of this study is that we did not examine micro-or long non-coding RNAs (77, 78). A second limitation is that our subjects had multiple different KS tumor types, and the inter-individual differences in scRNAseq between and within tumors remains to be characterized. In addition, subjects with epKS had received differing durations of antiretroviral therapy. Nevertheless, the current study demonstrates the feasibility and utility of scRNAseq for interrogating the KS biology and identifying potential prognostic and predictive biomarkers.

Taken together, these studies demonstrate the feasibility of single-cell, multiomic analyses to characterize the malignant and stromal composition of primary KS blood and tumor tissue, quantitate viral and host gene expression, identify prognostic and predictive biomarkers and potential therapeutic targets, and evaluate the efficacy of therapeutic interventions.

## Materials and Methods

### Biopsies and skin cell dissociation

Viably frozen in (Bambanker, serum-free cell freezing media. Sigma) single-cell suspensions were prepared from fresh primary skin biopsy samples using enzymatic digestion and gentle manual tissue dissociation (Whole Skin Dissociation Kit, Miltenyi Biotech) and thawed immediately prior to submission for single cell RNA sequencing. It should be noted that sample preparation without enzymatic digestion dramatically diminished the diversity and abundance of cell populations obtained from skin lesions.

### scRNAseq

Libraries were prepared using the 10x Genomics 5′ immune profiling kit-snRNA-seq protocol (GTAC@MGI). The resulting 10x library was sequenced on an Illumina S4 flow cell (300 cycles targeting 100,000 reads/cell). Alignment and gene expression quantification were performed with CellRanger multi pipeline (v7.1.0). The feature-barcode matrices were QCed, normalized and scaled using Seurat’s (v4.2.1) default settings. Principal component (PC) analysis was performed based on selected high variable genes and clustering of cells was performed using resolution = 0.7. Dimensionality reduction and visualization were performed using Seurat’s tSNE and UMAP functions. Cells were annotated with SingleR (v2) using expression profiles from the Human Primary Cell Atlas (HPCA) dataset. Finally, differential expression analyses was performed using Seurat’s FindMarkers function with the Wilcoxon Rank Sum method (logfc.threshold=0.2, min.pct=0.05). Volcano plots were generated using EnhancedVolcano (v1.16.0) to visualization of the top differentially expressed genes.

Reads from single cells obtained for each sample were mapped against the human genome, the KSHV/HHV8 genome, and the HIV-1 genome, visualized using t-distributed stochastic neighbor embedding (t-SNE) or uniform manifold approximation and projection (UMAP) plots, and clustered to enable cell type identification. Analysis: Default parameters for t-SNE clustering used the top 10 principal components from the principle component analysis (PCA) step as initialization for secondary analysis. The reference dataset from the Human Primary Cell Atlas (HPCA) was used to annotate clusters using SingleR.

### ddPCR

Viably frozen single-cell suspensions of primary KS skin biopsy samples were thawed and DNA was extracted from isolated cells using DNeasy kit (Qiagen). Then, digital droplet PCR was used to quantify copies of viral gene K1 using the QX200 Droplet Digital PCR System (BioRad). Probe sequence was 5’-/56-FAM/CGG CCC TTG /ZEN/TGT AAA CCT GTC /3IABkFQ/ -3’, and the primer sequences were 5’-GTT CTG CCA GGC ATA GTC -3’ and 5’-GCC AGA CTG CAA ACA ACA TA -3’. Results were detected with QX200 Droplet Reader (BioRad).

## Acknowledgements

This work was supported by Public Health Service grants to L.R. (R21 CA257493), supplemental funding from the Siteman Cancer Center (P30 CA091842) and the AIDS Malignancy Consortium (U01 CA121947) . We thank the Alvin J. Siteman Cancer Center at Washington University School of Medicine and Barnes-Jewish Hospital in St. Louis, MO. and the Institute of Clinical and Translational Sciences (ICTS) at Washington University in St. Louis, for the use of the Genome Technology Access Center, which provided scRNAseq services and support. The Siteman Cancer Center is supported in part by an NCI Cancer Center Support Grant CA091842 and the ICTS is funded by the National Institutes of Health’s NCATS Clinical and Translational Science Award (CTSA) program grant #UL1 TR002345.

## Financial Statement

The funders had no role in study design, data collection and analysis, decision to publish, or preparation of the manuscript.

